# Radiation Induced Metabolic Alterations Associate with Tumor Aggressiveness and Poor Outcome in Glioblastoma

**DOI:** 10.1101/2020.02.04.933648

**Authors:** Kshama Gupta, Ivan Vuckovic, Song Zhang, Yuning Xiong, Brett L. Carlson, Joshua Jacobs, Ian Olson, Xuan-Mai Petterson, Slobodan I. Macura, Jann Sarkaria, Terry C. Burns

## Abstract

Glioblastoma (GBM) is uniformly fatal with a one year median survival rate, despite the best available treatment, including radiotherapy (RT). Impacts of prior RT on tumor recurrence are poorly understood but may increase tumor aggressiveness. Metabolic changes have been investigated in radiation-induced brain injury; however, the tumor-promoting effect following prior radiation is lacking. Since RT is vital to GBM management, we quantified the tumor-promoting effects of prior radiotherapy (RT) on patient-derived intracranial GBM xenografts and characterized the metabolic alterations associated with the protumorigenic microenvironment. Human xenografts (GBM143) were implanted into nude mice 24h following 20Gy cranial radiation vs. sham animals. Tumors in pre-radiated mice were more proliferative and more infiltrative, yielding faster mortality (p<0.0001). Histologic evaluation of tumor associated macrophage/microglia (TAMs) revealed cells with a more fully activated ameboid morphology in pre-radiated animals. Microdialyzates from the radiated brain at the margin of tumor infiltration contralateral to the site of implantation were analyzed by unsupervised liquid chromatography-mass spectrometry (LC-MS). In pre-radiated animals, metabolites known to be associated with tumor progression (like, modified nucleotides and polyols) were identified. Whole-tissue metabolomic analysis of the pre-radiated brain microenvironment for metabolic alterations in a separate cohort of nude mice using ^1^H-NMR revealed significant decrease in levels of antioxidants (glutathione (GSH) and ascorbate (ASC)), NAD+, TCA intermediates, and increased ATP and GTP. Glutathione and ASC showed highest VIPpred (1.65) in OPLS-DA analysis. Involvement of ASC catabolism was further confirmed by GC-MS. To assess longevity of radiation effects, we compared survival with implantation occuring 2 months vs. 24h following radiation, finding worse survival in animals implanted at 2 months. These radiation-induced alterations are consistent with a chronic disease-like microenvironment characterized by reduced levels of antioxidants and NAD+, as well as elevated extracellular ATP and GTP serving as chemoattractants, promoting cell motility and vesicular secretion with decreased levels of GSH and ASC exacerbating oxidative stress. Taken together, these data suggest that IR induces tumor-permissive changes in the microenvironment with metabolomic alterations that may facilitate tumor aggressiveness with important implications for recurrent glioblastoma. Harnessing these metabolomic insights may provide opportunities to attenuate RT-associated aggressiveness of recurrent GBM.

## 1. Introduction

Glioblastoma multiforme (GBM; WHO grade IV) is the most common adult primary brain malignancy (1,2), accounting for 50% of all gliomas across all age groups (2). Standard treatment includes surgical resection, radiation therapy (RT), and chemotherapy; however, the overall five-year survival rate is <10% with mortality approaching 100% (3,4) is unfavorable prognosis may be due to the high propensity of tumor recurrence, with many recurring within one year, and 90% of these tumors forming within the prior RT field (5–7).

Radiation-induced changes in the brain and tumor microenvironment injury resulting in molecular, cellular, and functional changes that can facilitate tumor aggressiveness upon recurrence(8). Such changes include decreased vascularity, innate immune activation, and altered pharmacokinetics, pharmacodynamics, and therapeutic efficacy of chemotherapy agents (9–12). Additionally, irradiation (IR) generated reactive oxygen and nitrogen species (ROS/NOS) play havoc with cellular proteins, DNA and phospholipid membrane (13). Mitochondria exposed to radiation produce increased reactive oxygen species (ROS), that may contribute to RT-induced cell senescence (14–16).

Tumor cell metabolism is strikingly different from that of normal cells with a shift from energy-producing pathways to those generating macromolecules necessary for proliferation and tumor growth. Through a tricarboxylic acid cycle (TCA), healthy cells metabolize glucose and produce carbon dioxide within an oxygen-rich environment, which efficiently produces a large quantity of adenosine triphosphate (ATP) (17). In hypoxic environments, these cells produce large quantities of lactic acid by anaerobic glycolysis. Conversely, in aerobic conditions, tumor cells rely on glycolysis for energy production (18), resulting in elevated rates of glucose uptake and increase lactate production (19). Lactate production during active tumor growth alters the tumor microenvironment (TME) by promoting acidosis, serving as a metabolic cancer cell fuel source, and inducing immunosuppression. RT may also have immunosuppressive effects leading to increased tumor aggressiveness, with associated increases in proliferation and infiltration (20), which may be exacerbated by prior RT.

Metabolic alterations may be pro-tumorigenic, promoting glioma initiation and progression (21–24). RT-induced metabolic changes in GBM depend on tumor volume, location, and dose-time regime of RT-administration, all of which can vary treatment response (8,25–29). While differential metabolism of glioma tumor cells can be targeted to regress tumor growth, understanding the impact of radiation-induced metabolic alterations in GBM microenvironment can provide new avenues to maximize long term benefits of RT in GBM care. The major objective of this study is to investigate the interactions between irradiation, tumor aggressiveness, and the associated metabolic changes in the tumor microenvironment. We quantify the tumor-promoting effects of prior radiotherapy (RT) on patient-derived intracranial GBM xenografts in mice and characterize the metabolic alterations associated with the protumorigenic microenvironment.

## 2. Materials and Methods

### 2.1. Ethics statement on mice

Six to eight weeks old female heterozygous Hsd: Athymic Nude-Foxn1nu/Foxn1+ mice were purchased from Envigo (Indianapolis, IN). Six to eight-week-old male C57BL/6J mice were purchased from Jackson Laboratories (Bar Harbor, ME). Mice were housed at the Mayo Clinic animal care facility, which is Association for Assessment and Accreditation of Laboratory and Animal Care International (AAALACI)-accredited. Aging was induced in two separate cohorts of C57BL/6J mice [fed with regular diet or high-fat diet (D12492, Research diets)] by keeping them in-housed for 24 months at the Mayo Clinic animal care facility, i.e. a small cohort of 5 mice (2 months old) was maintained for 22months fed throughout on regular diet to obtain an aged mice group (24month old), and, another cohort of 5mice (2 months old) was fed on high-fat-diet (HFD) to induce obesity and continued on HFD for 22 months to obtain an aged-obese mice group (24month old). All animal procedures were performed with proper animal handling, adhering to the National Institutes of Health (NIH) guidelines and protocols approved by the Institutional Animal Care and Use Committee (IACUC) at Mayo Clinic, Rochester.

### 2.2. Cranial irradiation of mice

Cranial irradiation was administered using the X-RAD SmART irradiator (Precision X-ray, North Branford, CT), which uses a cone beam CT (CBCT) for accurate target localization. The stereotactic coordinates were determined from the target-set on CBCT using the first scan for each mouse within all groups (values ranged between x = 0.25 to 0.35, y = −3.8 to −4.0, and z = −5.8 to −5.95, depending on mice and strain-type). Whole brain radiotherapy was performed as described (30), using parallel opposed lateral beams with 10mm square collimator. Radiation treatments included 10, 15, or 20Gy single dose [20Gy] administration, or 4Gyx10 dose-fractionation. Control group mice were handled similarly as the treated, but with no radiation dose administered (0Gy).

### 2.3. Intracranial injections in mice

Intracranial injections in athymic nude mice were performed as previously described (31). Briefly, GBM143 cells were obtained from flank tumors and cultured *in vitro*. These cultured cells were dissociated using TryplE (Cat# 12563011, Thermo Scientific) and resuspended in PBS at a concentration of 100,000 cells/μl (with injection volume 3μl/mouse). Mice were anesthetized using Ketamine: Xylazine mixture (100mg/kg Ketamine and 10mg/kg Xylazine) injected intraperitoneally (IP) with a 0.5cc syringe. The surgical procedure involved the following steps: disinfecting mice head with Betadine, lubricating the eyes with artificial tears, making a 1cm midline incision extending from just behind the eyes to the level of the ears using sterile scalpel while applying pressure to have the incision open. Using a cotton swab, the skull was cleared to have the bregma exposed, a point 1mm anterior and 2mm lateral from bregma was identified and drilled through the skull using an 8bit Dremel drill. For stereotactic injection, Hamilton syringe with a 26G needle assembly was cleaned thoroughly, fixed on the injection jig, and 3μl of cell suspension drawn into it. Injection jig was sterilized by wiping with STERIS Spor-Klenz and draping it with a sterile towel. The mouse having its skull drilled was placed on the jig and fixed using front teeth hook at mouthpiece and ear pins. Using the stereotactic controls, the needle was inserted 3mm deep into the brain and, 300,000cells/3μL were injected at a rate of 1μL/min for over 3min using the syringe pump. The needle was maintained as inserted in place inside the skull for additional 1min, and then drawn out gently using the stereotactic controls. Mouse skull hole using sealed using bone cement, and the wound sutured with 4-0 vicryl with rb-1 needle (Ethicon J304H). Triple antibiotic was applied to the incision and stitches to prevent infection, and the mouse was left in the warm cage to recover from anesthesia. Water was supplemented with children’s ibuprofen starting 24hrs prior to starting the procedure and continued for 48hrs post-surgery.

### 2.4. Histology and immunofluorescence

Athymic nude mice injected with the established PDX line, GBM143, were euthanized using isoflurane overdose at day of moribund (i.e. after 58days of tumor cell-implantation). PBS cardiac perfusion was performed prior to termination under fully anesthetized conditions to remove the circulating peripheral leukocytes from the brain. Brains were extracted, fixed in 10% buffered formalin for 24hours, paraffin embedded, and 5μm coronal sections were obtained. All processing after fixation was performed at Mayo Clinic Histology core, Scottsdale. For histologic analysis, slides were stained with hematoxylin and eosin (H&E) and visualized by bright field microscopy at 4X microscopic magnification using Leica DMI-6000B (software: Leica Application Suite X (Leica Microsystems, Wetzlar, Germany)). Percent positive HE stained area was assessed to estimate relative tumor burden between the samples.

HE stained sections were reviewed to identify appropriate tumor bearing regions and respective unstained slides processed for immunofluorescence staining with human Lamin A+C and Ki67 antibodies using standard procedure. Briefly, slides were deparaffinized in xylene and rehydrated by washing (3mins each) in serially diluted ethanol from 100%, 95%, 75%, 50%, and then distilled H_2_O. Antigen retrieval was performed using pre-warmed 9.8mM Sodium citrate buffer (pH=6.0, with 0.05%Tween 20) for 30mins in hot steamer. Slides were rinsed in distilled H_2_O and PBS, blocked in blocking solution (10% N goat serum and 1% BSA in PBS), and stained with primary antibody (diluted in blocking solution, 1:300) for overnight in humidified chamber at 4°C. The slides were washed in PBS (3×5mins), stained with secondary antibody (diluted in blocking solution, 1:300) for 2hrs at room temperature, washed, and mounted with ProLong Gold reagent having DAPI (P36935, Life Technologies). Images were acquired at 4X microscopic magnification and tiling done using Leica DMI-6000B (software: Leica Application Suite X).

#### 2.4.1. Image analysis

All IF stained slides were quantified and scored for single cell count in a defined region with x-y coordinates approximated at tumor center (for Lamin A+C, and Ki67) or at centre of corpus callosum (for Lamin A+C), respectively, using Image J (32,33) and CellProfiler 2.2.0 (Broad Institute of Harvard and MIT) (34). Briefly, for single cell counting, an IF image obtained was imported into Image J, threshold was set, channels split, and image in relevant single channel was selected and converted to black and white (BW). An area template having fixed size was generated to define a contained region at tumor or at the center of corpus callosum, respectively, maintaining consistency between different sample slides. This defined area selectively masked was overlaid and appropriately positioned in the BW image, and all background cells out of the masked region were eliminated. The resultant image was transferred to CellProfiler 2.2.0 software (Broad Institute of Harvard and MIT) (34), the masked region was cropped and used as input; the pipeline for single cell counting was run to detect nuclei and quantify cells within this defined region.

To evaluate microglial activation, slides were stained for Iba-1 using standard procedure for IF. Images represented with 20X magnification were acquired on Leica DMI-6000B (software: Leica Application Suite X) and 40X magnification on Zeiss Axio Observer Z.1 (Software: Zen 2.3 SP1, Jena, Germany). Microglial morphology was assessed using ImageJ (32,33). Antibodies used: Rabbit monoclonal Anti-h-Lamin A+C [EPR4100] (Cat# Ab108595, Abcam, Cambridge, United Kingdom); Rat monoclonal Ki67 (SolA15) (Cat #14-5698-82, eBioscience Invitrogen, Waltham, MA); Rabbit monoclonal Anti-Iba-1 (Cat# 019-19741, Wako). Secondary antibodies from Jackson ImmunoResearch Laboratories, Inc. (West Grove, PA) included polyclonal affinity-pure whole IgG:Cy3-Goat Anti-Rabbit IgG (H+L) (code: 111-165-003) and Cy5-Goat Anti-Rat IgG (H+L) (code: 112-175-143).

### 2.5. Microdialysis

To evaluate changes in the extracellular milieu of radiated brain, a small group of mice (3 mice per group) from 0Gy and 20Gy single-dose irradiated mice cohorts injected with GBM143 24hr post-IR, were microdialyzed on their contralateral hemisphere (non-tumor bearing side) at day 30 from tumor cell injection. The microdialysis set-up and surgical procedure was followed as described from the facility of Dr. Doo-Sup Choi, at Mayo Clinic (35). Briefly, the mice were housed singly for 2hrs in the microdialysis room to acclimatize, and then anesthetized using Ketamine:Xylazine mixture. Survival surgery was performed on a rotating platform with stereotactic guidance under sterile conditions. A microdialysis probe with a 2.0mm cellulose membrane (Brain Microdialysis, CX-I Series, Eicom, Kyoto, Japan; MW cut off: 50,000 Da) was inserted at a point 1mm anterior and 2mm lateral from bregma on the contralateral hemisphere and secured to the guide cannula. The probe was connected to a microsyringe pump (Eicom, Kyoto, Japan), which delivered Ringer’s solution (145 mM NaCl, 2.7 mM KCl, 1.2 mM CaCl_2_, 1.0 mM MgCl_2_, pH 7.4) at a 1.0 μl/min flow rate. The samples were collected in 0.2ml collection tubes maintained at 4°C for 3.5 hrs, and then immediately frozen and stored at −80°C until analyzed.

### 2.6. Metabolomics

#### 2.6.1. Proton Nuclear Magnetic Resonance spectroscopy (^1^H-NMR)

Athymic nude mice, 0Gy-control, and 20Gy single-dose irradiated (10 mice per group) were sacrificed and immediately frozen in liquid nitrogen. Brian tissues were collected on dry ice and pulverized in liquid nitrogen. The pulverized mouse brain tissue (~55-60 mg) was homogenized and extracted with 300μl of ice-cold 0.6M perchloric acid (HClO_4_) solution. Sample tubes were (36)extraction procedure was repeated on the pellets (with ~150μL HClO_4_) and supernatant obtained from two rounds of extraction were combined and neutralized with 140μl of 2M potassium bicarbonate, KHCO_3_. In 400μL aliquot of neutralized extract, 100μL of 0.1M phosphate buffer and 50μL of 1mM TSP-d4 in D_2_O were added. Samples were vortexed for 20 seconds and transferred to 5mm NMR tubes. The NMR signal was acquired on Bruker AVANCE III 600 MHz instrument (Bruker, Billerica, USA). ^1^H-NMR spectra were recorded using 1D NOESY pulse sequence with presaturation (noesygppr1d) under the following conditions: 90-degree pulse for excitation, acquisition time 3.90s, and relaxation delay 5s. All spectra were acquired with 256 scans at room temperature (298K) with 64k data points and 8417Hz (14 ppm) spectral width. The recorded ^1^H-NMR spectra were phase and baseline corrected using TopSpin 3.5 software (Bruker, Billerica, MA). The spectra were then processed using Chenomx NMR Suite 8.3 software (Chenomx Inc., Edmonton, Canada). The compounds were identified by comparing spectra to database Chenomx 600MHz Version 10 (Chenomx Inc., Edmonton, Canada) and literature data (36–42). Quantification was based on an internal standard (TSP-d4) peak integral. The metabolite concentrations were exported as μM in NMR sample and recalculated as μmol/g of wet tissue.

#### 2.6.2. Gas chromatography–mass spectrometry (GC-MS)

For GC-MS analysis, 70μl neutralized brain extracts (approx. 6.4mg of tissue wet weight) from athymic nudes were obtained using perchloric acid extraction method with 2M KHCO_3_ based neutralization as described for 1H-NMR, centrifuged at 10,000g for 10mins, and cleared supernatant collected in fresh 1.5ml eppendorf tubes. These samples were completely dried in a SpeedVac concentrator run overnight. They were subsequently methoximated using 10μL MOXTM reagent (Cat# TS-45950, ThermoScientific, Waltham, MA) at 30°C for 90min and then derivatized using 40μL of N-methyl-N-trimethylsilyl trifluoroacetamide with 1% trimethylchlorosilane (MSTFA+1% TMCS: Cat# TS48915, ThermoScientific, Waltham, MA) at 37°C for 30min. Metabolite levels were determined using GC-MS (Hewlett-Packard, HP 5980B) with DB5-MS column. GC-MS spectra were deconvoluted using AMDIS software (NIST, Gaithersburg, MD) and SpectConnect software (Georgia Tech, Atlanta, GA, USA) was used to create metabolite peaks matrix. The Agilent Fiehn GC/MS Metabolomics RTL Library (Agilent, Santa Clara, CA) was used for metabolite identification. Ion count peak area was used for analysis of the relative abundance of the metabolites (43).

Similar to above, whole brain extracts using perchloric acid method were also prepared from a cohort of C57BL/6J mice and evaluated by ^1^H-NMR and GC-MS. C57BL/6J mice included in the study were divided into 5 groups (with 4-5 mice per group) as follows: control (0Gy), 20Gy single-dose irradiated, 4Gyx10 fractionation-dose irradiated, and, two aged (24mo) non-irradiated mice groups, aged (24mo): fed on regular diet and, aged-obese (24mo): induced with obesity using high-fat-diet. All mice otherwise were 5-6 months old.

#### 2.6.3. Data analysis

Multivariate analysis of NMR data was performed using SIMCA 15 software (Sartorius Stedim Biotech, Göttingen, Germany). Principal component analysis (PCA) was used to detect any innate trends and potential outliers within the data. Supervised Partial Least Squares discriminant analysis (PLS-DA) and Orthogonal Projections to Latent Structures Discriminant Analysis (OPLS-DA) were performed to obtain additional information including differences in the metabolite composition of groups, variable importance in the projection (VIP) values, and regression coefficients. OPLS-DA models were calculated with unit variance scaling and the results were visualized in the form of score plots to show the group clusters. The VIP values and regression coefficients were calculated to identify the most important molecular variables for the clustering of specific groups. Nonparametric Wilcoxon rank sum test and Student T-test were performed to determine the statistically significant differences between the groups with significance considered at level ≤0.05.

### 2.7. Survival curves

Athymic nudes, grouped as control (non-irradiated, 0Gy) and irradiated with 20Gy single dose, were divided into two study cohorts: 1) Short-term IR: where 5 mice from each group were injected with GBM143 cells after short-term prior IR-exposure of 24hrs and 2) Long-term IR: where 5 mice from each group were maintained for 2months post-irradiation and then injected with GBM143 cells. Survival time (in days) for each mouse was recorded until 70days post tumor cell injection. The overall survival was calculated by Kaplan-Meier method and log-rank test was used to compare the survival curves (44). A p-value of ≤0.05 was considered to be statistically significant.

### 2.8. Statistical representation

The difference between specific metabolites or a parameter measured across two groups was estimated for p-value, q-value or False discovery rate (FDR), and Fold change (FC), as appropriate. Graphs were plotted using software(s): GraphPad Prism 8.2.0 (GraphPad, San Diego, CA), Heatmapper (Wishart Research Group, University of Alberta and Genome Canada) (45) and Microsoft Office Excel. Statistical significance is represented as p-values: *p≤0.05; **p≤0.01; ***p≤0.001, ****p≤0.0001, or, q-values, where specified.

## 3. Results

### 3.1 Effect of radiation on tumor growth, proliferation and migration

The experimental strategy is shown in Figure 1 and Supplementary Figure 2. Mice were cranially irradiated with either 20Gy (single dose) or 0Gy (control). At 24hrs post-irradiation, a pre-established GBM line (GBM143) was injected into the mice brain. Tissues were collected at moribund and evaluated with histology for tumor growth. A small cohort of mice radiated with 10Gy (single dose) and injected with GBM143 line was also compared with the 0Gy and 20Gy cohorts for relative tumor burden using haematoxylin and eosin (H&E) staining. No difference in tumor size was observed between 0Gy and 10Gy; however, 20Gy irradiated samples had significantly higher percent of section area positive for tumor by H&E, indicating an overall faster rate of tumor growth (Supplementary Figure 1). Thus, 10Gy cohort was not pursued for further evaluation. Sections from 20Gy and 0Gy were analyzed for tumor growth and proliferation using human-Lamin A+C and Ki67 staining. Significantly larger tumors were observed in the 20Gy group. More h-Lamin A+C cells were seen in the corpus callosum of 20Gy mice (Figure 1B). Quantitative analysis was performed by counting both h-Lamin A+C and Ki67 within the tumor to evaluate proliferation. Similarly, h-Lamin A+C was assessed in the midline corpus callosum to evaluate cell migration towards the contralateral hemisphere, as illustrated in Figure 1C. Significantly higher numbers of h-Lamin A+C and Ki67 cells were seen in 20Gy mice, as compared to 0Gy control at tumor. Similarly, significantly higher h-Lamin A+C staining was observed in 20Gy compared to 0Gy mice in the midline corpus callosum, suggesting a higher number of cells migrating toward the contralateral hemisphere (Figure 1D).

**Figure 1:**
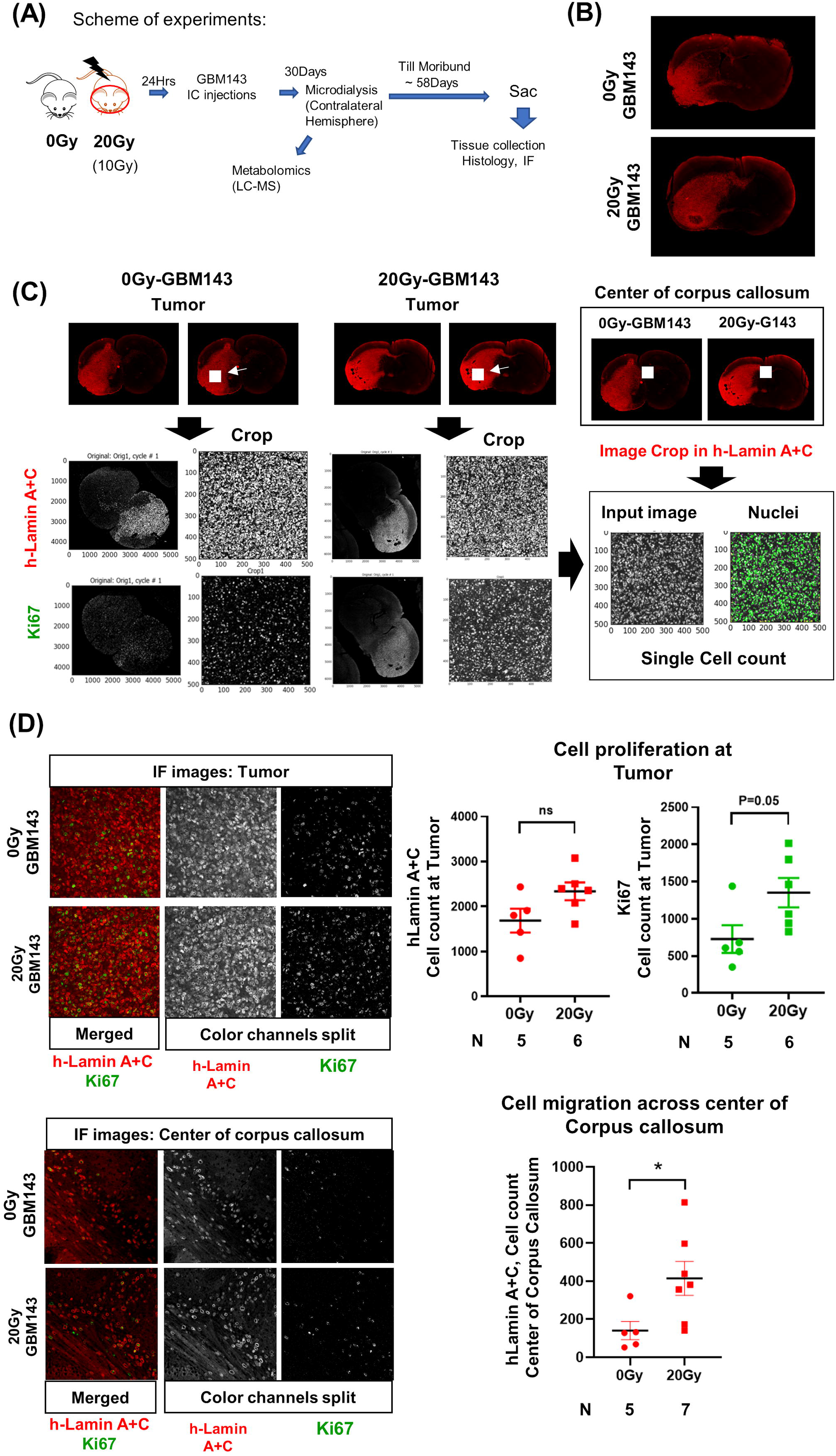
A, Scheme of experiments. B, Representative IF images (at 4X, tiling) for hLamin A+C staining from 0Gy and 20Gy-IR mice coronal sections to assess tumor growth and invasion. C, Scheme illustrating steps involved in performing single cell count: mice brain coronal sections are stained for hLamin A+C ‒Cy3 (and Ki67 – Cy5), for both 0Gy and 20Gy. A defined region is selected and masked (area-squared in white). This masked area-image in single channels is imported into cell profiler software and cropped. This cropped image is used as the input image, pipeline for nuclei detection run, and single cell count obtained. Similar steps are performed for a defined region selected at centre of corpus callosum for hLaminA+C staining (images in box, on right). D, Top-panel: Representative images (at 20X) show IF staining at tumor; and the dot-plot of single cell count for hLamin A+C and Ki67 staining. Bottom-panel: Representative images (at 20X) show IF staining at center of corpus callosum, and dot-plot of single cell count for hLamin A+C. Statistical significance is represented as *p ≤ 0.05; **p ≤ 0.01.

**Figure 2:**
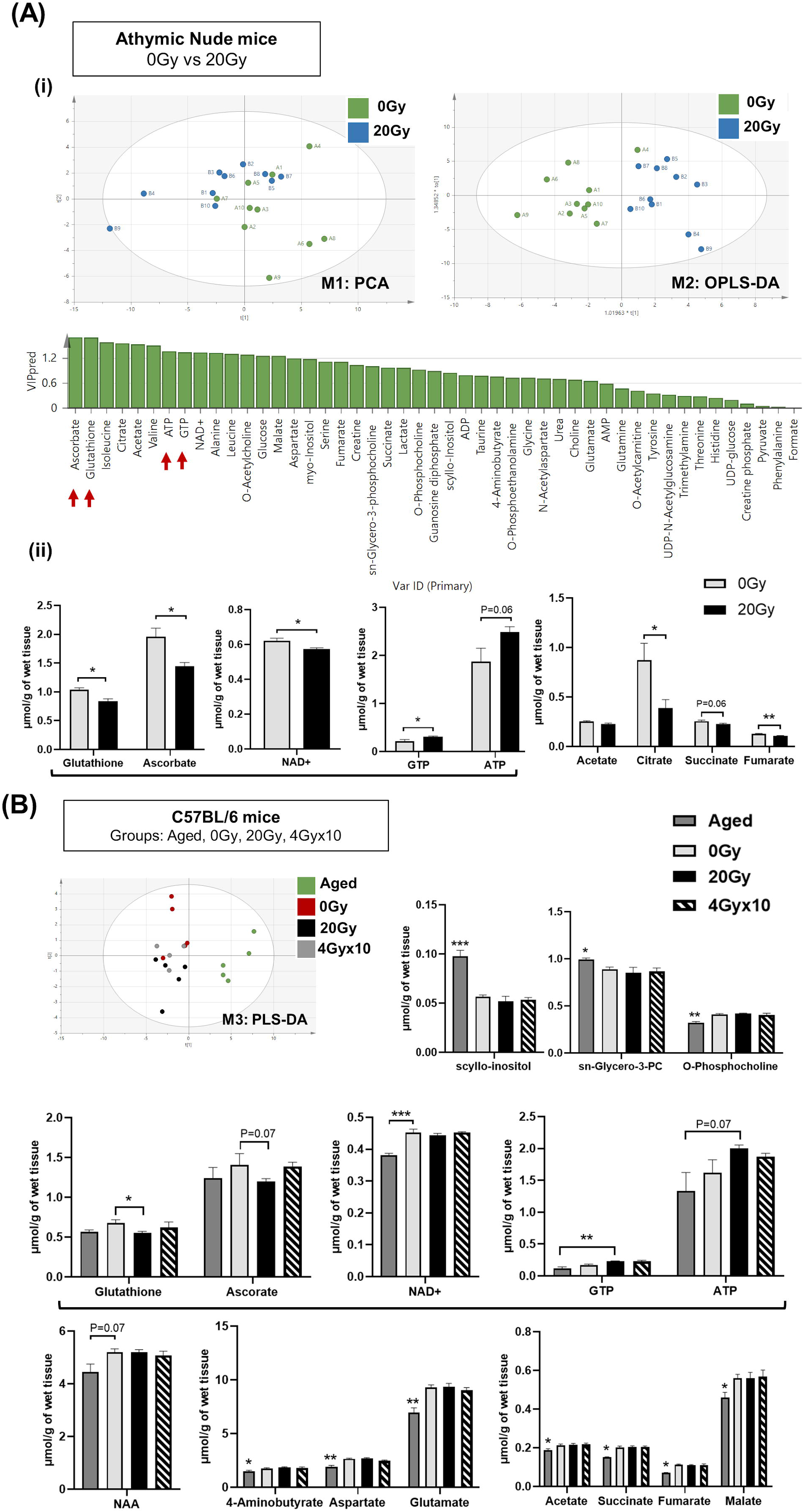
Proton nuclear magnetic resonance (^1^**H NMR**): A, (i) Principal component analysis, PCA between 0Gy and 20Gy. Supervised Orthogonal Projections to Latent Structures Discriminant Analysis, OPLS-DA shows further separation of 20Gy mice group from non-irradiated control based on differences in the metabolite composition of groups with predicted-VIP (variable importance in the projection) values shown in graph below. (ii) The graphs show, significantly altered metabolites between the 0Gy vs 20Gy. B, Multivariate analysis of NMR data performed using SIMCA 15 software for cohort of C57BL/6 mice, having groups as indicated. Supervised Partial Least Squares discriminant analysis (PLS-DA) performed shows, separation of all the four groups. Bar graphs show metabolites most significantly altered between groups. Statistical significance is represented as *q ≤ 0.05; **q ≤ 0.01.

### 3.2 Metabolomics

#### 3.2.1 Microdialysis

To assess for radiation-induced changes in the extracellular milieu, a pilot experiment with intracranial microdialysis was performed (using a microdialysis probe with 2.0mm cellulose membrane for Brain Microdialysis, CX-I Series, Eicom; MW cut-off: 50KDa) as described (35), in the contralateral hemisphere of a small cohort of mice from each study group (0Gy and 20Gy) at day 30 post-irradiation and GBM143 injection. Microdialysates were analyzed for untargeted liquid metabolic profiling using liquid chromatography-mass spectrometry (LC-MS), as described (46) (Methods described in Supplementary Materials). Principal component analysis could separate the groups 0Gy and 20Gy, indicating metabolic changes in effect of irradiation. A trend toward elevated levels of metabolites relevant to cancer progression was observed in the 20Gy mice, including modified nucleotides (N6-methyladenosine, pseudouridine), polyol (myo-inositol, quebrachitol) detected in the 20Gy (Supplementary *Excel* Sheet 3). However, there were very limited identifiable metabolites with a total of <60 due to low sample volume obtained after a 3.5hr microdialysis run at run rate of 1ul/min (Supplementary *Excel* Sheet 3). Moreover, due to technical challenges involved with keeping ≥4 mice per group in microdialysis and the limited volume of microdialysates collected for evaluation, significant conclusions could not be made. We therefore utilized a whole tissue metabolomics approach in non-tumor bearing mice to evaluate the metabolic changes post-irradiation.

#### 3.2.2 Proton nuclear magnetic resonance (^1^H NMR) spectroscopic analysis

We sought to identify the radiation induced metabolic alterations in the brain stroma associated with the observed outcome of higher tumor growth and proliferation in 20Gy mice. The scheme for the mice groups involved is illustrated in Supplementary Figure 2A (i). Whole brain metabolomics was performed in two separate mouse strains, athymic nude mice and C57BL/6 mice. Athymic nude mice were included since the tumor study described above was performed with human-PDX line in athymic nudes; C57BL/6 mice were included to eliminate strain dependence and to avoid potential confounding effects of immunodeficient mice. An aged-C57BL/6 mice (24 months old) with and without diet-induced-obesity were analyzed to assess whether or not the radiation-induced metabolic changes in the brain were similar to those induced by aging or obesity-induced senescence. A small group of C57BL/6 mice were administered a fractionated dose of 4Gyx10 for comparative analysis.

Data from proton nuclear magnetic resonance (^1^H NMR) spectroscopic analysis revealed clear separation of 0Gy and 20Gy mice cohorts from athymic nude mice, using PCA (Figure 2A(i)). Supervised Orthogonal Projections to Latent Structures Discriminant Analysis (OPLS-DA) further separated the two groups based on metabolite composition differences with predicted-variable importance in the projection (VIP) values shown. The most important molecular variables for clustering of specific groups include glutathione (GSH) and ascorbate (ASC) having VIPpred 1.65, along with differences in ATP and GTP levels as potentially distinguishing characteristics. After IR a significant reduction of GSH, ASC, and NAD^+^ levels were observed, along with increases in ATP and GTP. Additionally, an overall reduced trend in TCA intermediates was observed in 20Gy. The multivariate analysis of NMR data performed using SIMCA 15 software for C57BL/6 mice demonstrated separation of groups: Aged 24mo, Aged-Obese 24mo, Control (0Gy), 20Gy single-dose cranially irradiated, and 4Gyx10 cranial IR-fractionated (Supplementary Figure 2A (ii)). Supervised Partial Least Squares Discriminant Analysis (PLS-DA) showed separation of all five groups. Specifically, the aged-groups (aged:24mo and aged-obese:24mo) were separated into a different component compared to the 0Gy, 20Gy, and 4Gyx10 groups (Figure 2B and Supplementary Figure 2A (ii)). There was a better separation of groups shown in model:M4 (aged, 0Gy and 20Gy) as compared to those shown in model:M5 (0Gy, 20Gy and 4Gyx10) (Supplementary Figure 2 A(ii), Table 1 for model parameters). Comparing all irradiated mice (IR group: 20Gy and 4Gyx10 analyzed together) with 0Gy using PLS-DA and OPLS-DA showed significant group separation. The VIP-total and VIPpred value estimation indicated the metabolites most relevant to this group separation, which included GTP, ATP, GSH, and ASC (Supplementary Figure 2B).

The relative abundance of metabolites identified post-IR for 20Gy single dose from ^1^H NMR for C57BL/6 mice showed reduction in GSH and ASC levels and an increase in ATP and GTP (Figure 2B). No significant difference was observed between 20Gy and 4Gyx10. Alterations specific to the aged-group involved increased levels of scyllo-inositol and sn-glycero-3-phosphocholine with concomitant reduction in O-phosphocholine. Other metabolites reduced significantly in aged-mice, were NAA (N-acetyl aspartate), neurotransmitters, and intermediates of TCA cycle. List of metabolites detected for athymic nude mice and C57BL/6 by ^1^H NMR are included in Supplementary *Excel* sheet 1.

#### 3.2.3 Gas chromatography–mass spectrometry (GC-MS)

Lysates prepared by perchloric acid extraction for ^1^H NMR were further evaluated using Gas Chromatography–Mass Spectrometry (GC-MS). The heatmap for relative abundance of metabolites (i.e. changes in normalized total peak area for the metabolites) between athymic nude mice groups, 0Gy and 20Gy, cranial IR with single dose is illustrated in Figure 3A (i). While there was internal variation observed within these groups, only a few significantly altered metabolites in 20Gy were identified., which included an increased trend in urea and, a reduction in levels of creatinine (Crn), NAA (N-acetyl aspartate), and NAA/Crn ratio post-irradiation. Importantly, ascorbic acid was significantly reduced in 20Gy and threonic acid was increased, reflecting ascorbic acid catabolism. The heatmap for relative abundance of metabolites (i.e. changes in normalized total peak area for the metabolites averaged for all mice within each group) between the single dose cranially irradiated C57BL6 mice groups (Aged: 24mo, Aged-Obese: 24mo, Control (0Gy), and 20Gy) and fractionally irradiated 4Gyx10 is included in Figure 3B (ii). The clustering was performed using Heatmapper software, which clearly distinguished between aged mice and aged-obese mice from 20Gy and 4Gyx10 groups. The significantly altered metabolites between control (0Gy) and irradiated groups (20Gy and 40Gy-F) involved increased levels in urea but no significant change in Crn, N-acetyl aspartate (NAA), and NAA/Crn ratio. However, there was a consistent trend with reduced levels of ascorbic acid and increased levels of threonic acid observed post-irradiation. Collectively, the results of ^1^H-NMR and GC-MS, indicate involvement of ROS clearance with active utilization of GSH and ASC as antioxidants. Scheme for ASC and GSH cycle in clearance of reactive oxygen species (ROS) and the role of GSH in regeneration of ASC is illustrated, along with intermediates of ascorbic acid catabolism are represented in Figure 3C. Expected metabolic alterations upon irradiation involve an increase in levels of ROS, utilization and reduction in GSH and ASC, with concomitant increase in by-products of ASC catabolism, threonic acid (ThrO), and Oxalic acid (OxA) as indicated (Figure 3C).

**Figure 3:**
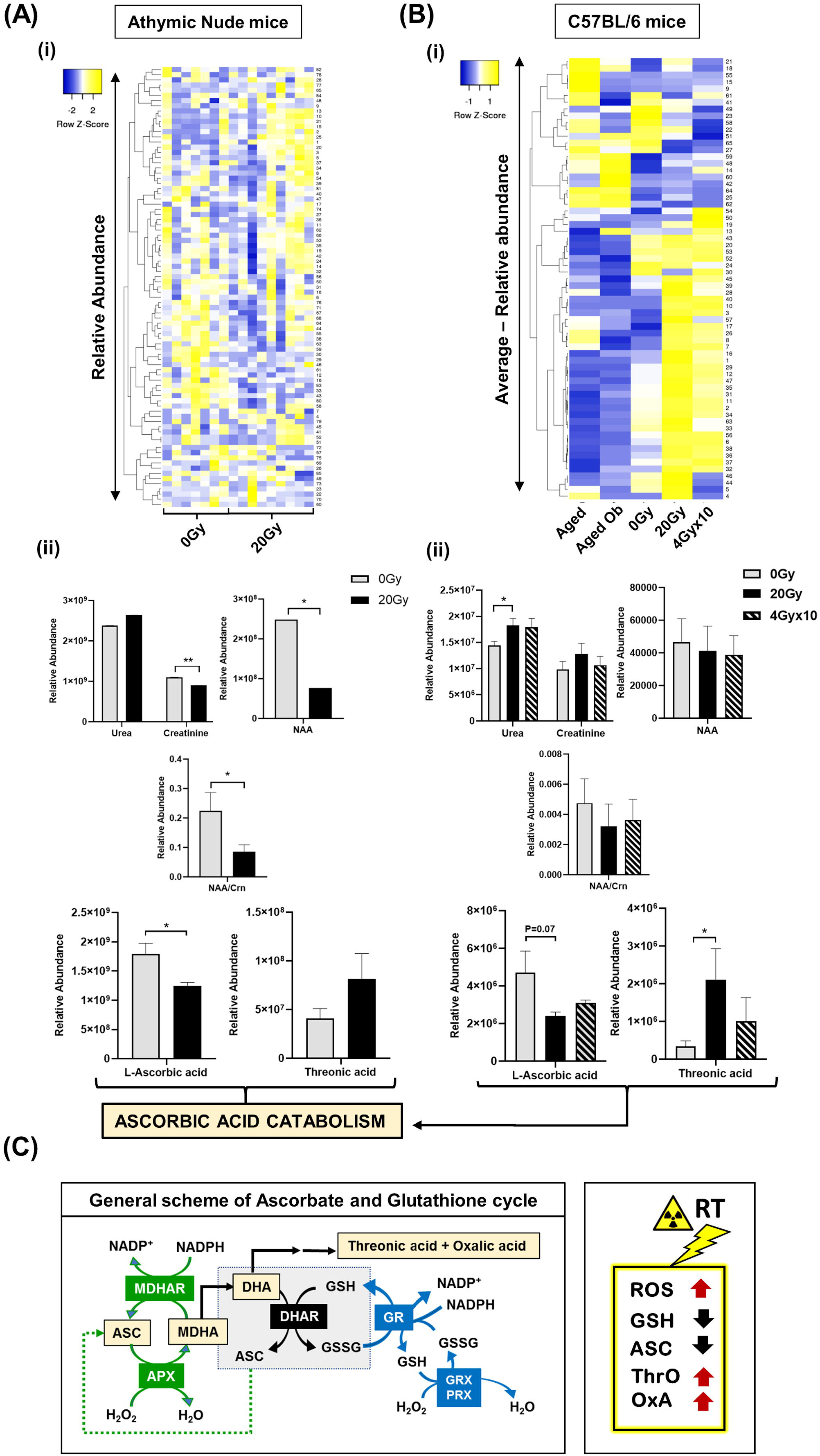
Gas chromatography–mass spectrometry (GC-MS): A, (i) Heatmap for relative abundance of metabolites (i.e. normalized total peak area of metabolites for all mice within each group) between athymic nude mice groups, 0Gy and 20Gy. (ii) The graphs show significantly altered metabolites between 0Gy and 20Gy mice. B, (i) Heatmap for relative abundance of metabolites averaged for each group (i.e. normalized total peak area for metabolites, averaged for all mice within each group), between C57BL/6 mice grouped indicated. (ii) The graphs show significantly altered metabolites between 0Gy, 20Gy and 4Gyx10. C, General scheme for ascorbate (ASC) and glutathione (GSH) cycle in clearance of reactive oxygen species (ROS). Intermediates of ascorbic acid catabolism are represented in orange boxes. Reactions in green show ASC-dependent peroxide metabolism; reactions in the central grey box show GSH-dependent regeneration of ASC; and reactions in red show GSH-dependent peroxide metabolism. To the right: illustrates the expected metabolic alterations upon irradiation, which include increases in levels of ROS, utilization of GSH and ASC, with concomitant increase in by-products of ASC catabolism, threonic acid, and Oxalic acid. ASC; MDHA, monodehydroascorbate; MDHAR, monodehydroascorbate reductase; APX, ASC peroxidase; GR, GSH reductase; GRX, Glutaredoxin; PRX, Peroxiredoxin; ThrO, L-threonic acid; OxA, oxalic acid; DHA, Dehydroascorbic acid, GSH, GSH reduced; GSSG, GSH oxidized; NADP^+^, nicotinamide adenine dinucleotide phosphate. Statistical significance is represented as *q ≤ 0.05; **q ≤ 0.01.

Other metabolites contributing to the separation of the groups in C57BL6 mice are shown in Supplementary Figure 3. There was no significant difference in levels of cholesterol in the aged-Obese group, which could be due to high internal variation observed within the groups or small cohort size of 5mice/group. However, there was a reduced trend in free fatty acids and overall higher cholesterol in group B, as compared to others. Notable metabolites separating the aged groups (aged-24m, and aged-Ob) from irradiated groups (20Gy and 4Gyx10) involved reduction in fumaric and succinic acids, and reduced levels of metabolic intermediates of glycolysis and Tricarboxylic Acid Cycle (TCA). Metabolic variations common to both aged and radiated mice cohorts included a rise in threonic acid, oxalic acid, D-allose, and myo-inositol. Additionally, there was a slightly higher trend in urea and Crn; however, this was not significant for either aged or irradiated mice groups. List of metabolites detected by GC-MS are included in Supplementary *Excel* sheet 2.

### 3.3 Immunostaining for microglia with Iba-1

To evaluate the inflammation caused by tissue irradiation and its crosstalk with injected GBM143 tumor cells, immunostaining was performed for microglia, with Iba-1 (Figure 4). Microglial morphology was assessed at 20X in ipsilateral (IH) and contralateral (CH) hemispheres for coronal slices from mice cerebral hemispheres, mice cranially irradiated with 0Gy and 20Gy-single dose, and injected 24hrs post-irradiation with GBM143 PDX line (Figure 4A, 4B). Microglia were observed to be enlarged, bushy, and branched for 0Gy-GBM143, as opposed to amoeboid for 20Gy-GBM143, indicating stages of higher activation and higher phagocytic activity for the 20Gy-GBM143 injected mice. To evaluate whether the observed microglial activation was a result of radiation alone or due to a combined effect of radiation with tumor injection, the microglial staining in ipsilateral hemispheres of 0Gy-GBM143 and 20Gy-GBM143 was compared with that of ipsilateral hemispheres of two separate mice cranially irradiated with 20Gy-single dose and not injected with any human-GBM PDX line. Intriguingly, negligible Iba1^+^ microglia staining was observed in the brain slices of 20Gy-IR alone, indicating the observed microglial activation to be an effect of crosstalk between irradiation and tumor pathogenesis. Figure 4D illustrates their relevance in our experimental setting with maximum microglial activation and phagocytic activity observed in 20Gy mice.

**Figure 4:**
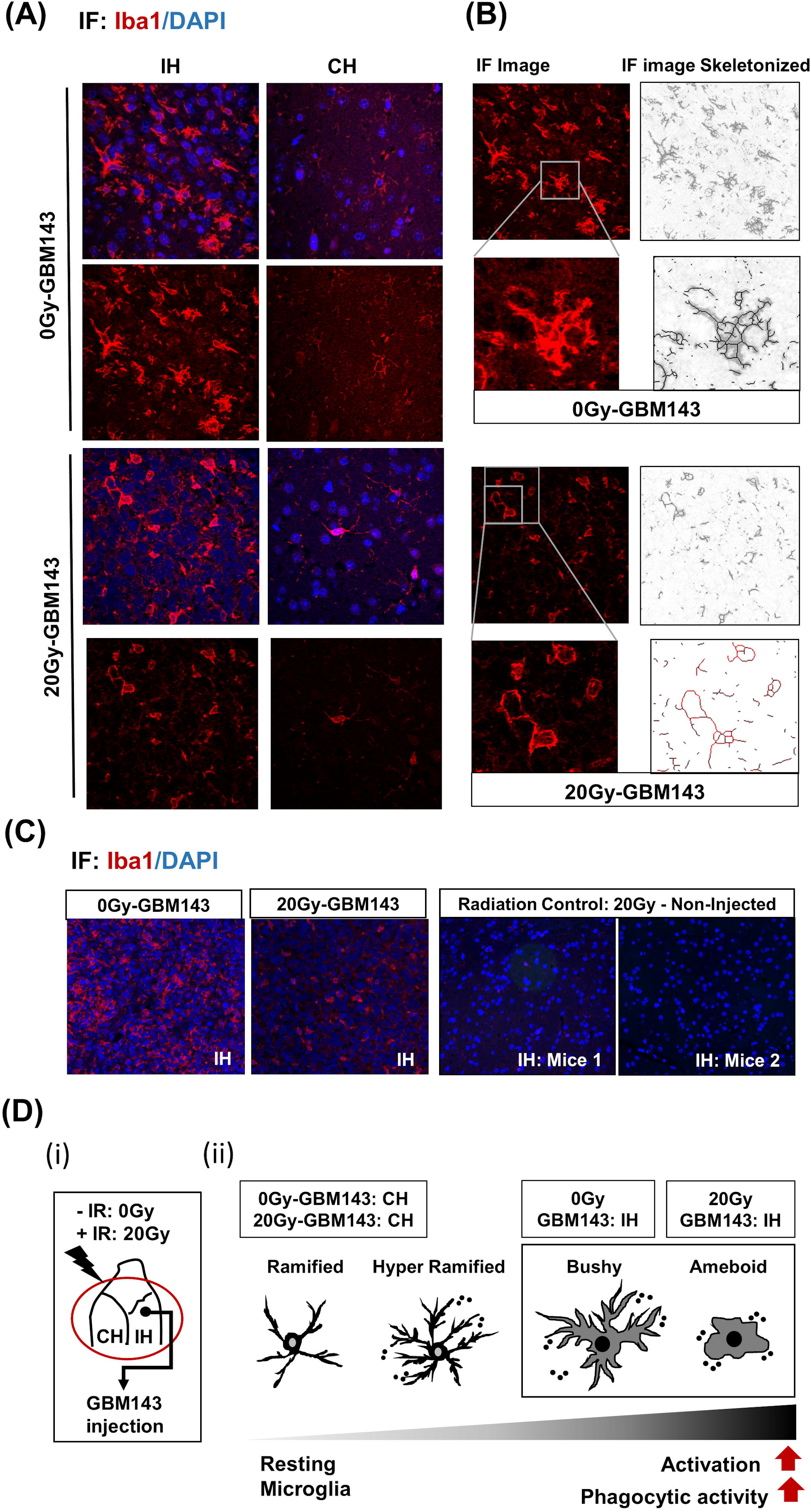
Immunostaining for microglia, with Iba-1: A, IF Images for microglial staining (Iba1/DAPI) and morphology (at 20X), in ipsilateral (IH) and contralateral (CH) hemispheres of 0Gy and 20Gy mice injected with GBM143 PDX line. B, the IF images skeletonized using Image J software to assess microglial morphology. C, the microglial staining in ipsilateral hemispheres of 0Gy-GBM143, and 20Gy-GBM143 compared with that of ipsilateral hemispheres of two separate mice cranially irradiated with 20Gy-single dose, however, not injected with any human-GBM PDX line. D, (i) Site of GBM143 injection at IH of mice having received ± cranial irradiation (ii) Stages of microglial activation observed in experimental setting.

### 3.4 Effect of radiation on GBM outcome

To further assess the effects of radiation-associated metabolic alteration on GBM outcome, survival analysis was performed in irradiated mice cohorts and their respective controls when injected with GBM143. To assess the short-term (ST-IR) or Long-term (LT-IR) effects of irradiation on GBM-outcome, the GBM143 cells were injected at two different time points: 1) 24hrs post-irradiation (ST-IR) or 2) two months post-irradiation (LT-IR). Significant reduction was seen in the survival of mice after irradiation ST-IR or, LT-IR (Figure 5). The combined graph of ST-IR and LT-IR further showed a significant difference is survival of 20Gy (ST-IR) and 20Gy (LT-IR), with median survival of 58 days and 51 days, respectively.

**Figure 5:**
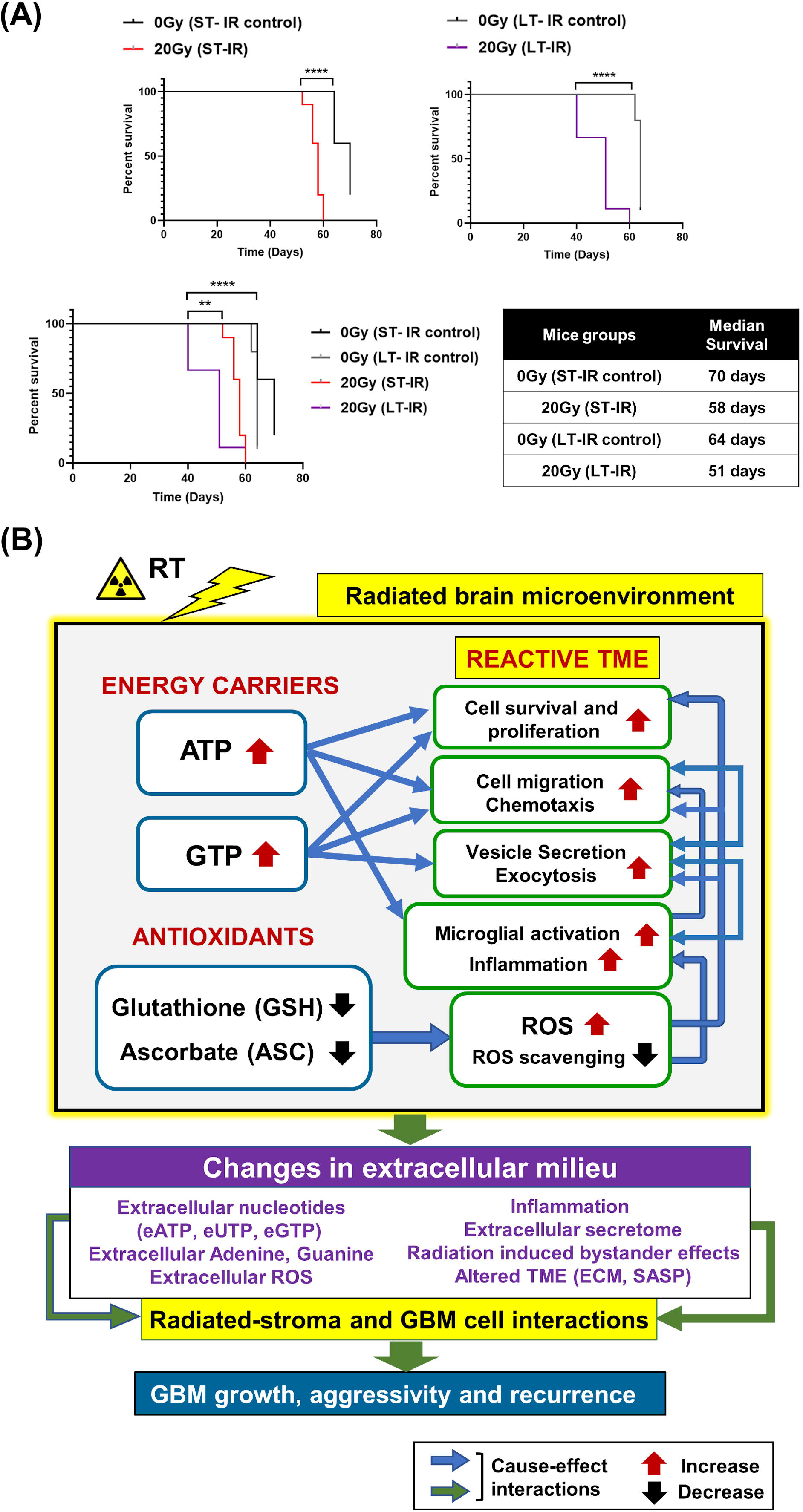
Effects of radiation-associated metabolic alteration on GBM outcome: A, Survival curves: Graphs show difference between survival of irradiated mice cohorts injected with GBM143 24hrs post-irradiation (ST-IR) or 2 months post-irradiation (LT-IR). B, Proposed model for radiation associated metabolic alterations and their effects on cell processes and GBM outcome. The box marked with yellow outline (irradiated by RT) shows metabolic changes in the radiated brain stromal microenvironment, with rise in energy carriers ATP and GTP, and reduction in levels of antioxidants, ascorbate and glutathione, and the cellular processes affected by them. With IR, continued and excess rise in levels of energy carriers and expense of antioxidants within stromal cells of the brain can lead to altered extracellular milieu. Pathophysiological changes in the extracellular milieu, which can be immediate or long term caused by the radiation are enlisted in purple box. These alterations would collectively contribute to radiated stroma and GBM cell interactions that are permissive to GBM growth, aggressivity and recurrence.

Collectively, our data demonstrate radiation-induced metabolic alterations, including a rise in energy carriers (ATP and GTP) and reduction in antioxidants (GSH and ASC) associated with tumor promoting cell processes (cell proliferation, migration and inflammation) and poor GBM outcome. The proposed model is illustrated in Figure 5B.

## 4. Discussion

This study aimed to identify the impact of high-dose radiation-induced brain injury on glioblastoma growth and aggressivity, and to probe the metabolic alterations in response to irradiation-associated growth. Radiation can severely impact the tumor microenvironment by altering the extracellular milieu at molecular and structural levels (8,25,47). Radiation treatment leads to production of ROS. Tumors adapt to oxidative stress through several mechanisms, including metabolic shifts and elevated antioxidant peptide production and intratumoral hypoxia generation (48–50). Metabolomics has emerged as the state-of-the-art approach to identify cancer cell state and biomarkers. Furthermore, metabolic profiling of tumor microenvironment can provide information on tumor cell fate (51–57). We evaluated the metabolic changes in the pre-irradiated brain microenvironment in response to 20Gy-IR and the association with observed tumor aggressivity and inflammatory microglial phenotype. Cell proliferation and migration are a direct function of the cell’s energy state (21); therefore, we involved ^1^HNMR to identify and quantify energy carriers as also described (21). Our data demonstrate elevated levels of ATP and GTP post 20Gy-IR, with reduced levels of antioxidants (glutathione and ascorbate) [Figure2]. This observation was conserved between both mice strains (athymic nudes and C57BL6) included in the study. Increased ROS levels are expected to occur in response to irradiation treatment, which can activate pro-tumorigenic signaling (21,22). Ascorbate and GSH serve as the prime cellular antioxidants. Glutathione can recycle itself and reduced ascorbate (58,59). A decline in relative abundance of ASC and concomitant rise in threonic acid was observed using GC-MS, which confirmed active ASC catabolism [Figure 3]. Decreased levels of ASC and GSH indicate active ROS scavenging. Their depletion due to increased demand can cause further accumulation of intracellular ROS.

ROS production is associated with DNA damage and cell death. However, chronically high levels of ROS in the tumor microenvironment can be pro-tumorigenic (50,60). Similarly, while ATP and GTP are essential components of cellular homeostasis, a rise in intracellular nucleotides (ATP and GTP) can cause their export out of the cell through extracellular vesicles, thus elevating extracellular levels of nucleoside and nucleotides (eATP, eGTP, adenosine and guanine) (61,62). Extracellular purinergic nucleotides can affect both stroma cell and tumor cell processes. Extracellular ATP has been implicated in facilitating chemotaxis of microglia, microglial activation, inflammation, and several neurological or neuropathological processes (63). Additionally, it can be internalized by tumor cells, increasing their intracellular ATP levels conferring metabolic reprogramming, increased tumor aggressivity, and treatment resistance (64–68). A recent lung cancer study has shown eATP to be involved in epithelial-to-mesenchymal transition, cell migration, and metastasis (69). While the biological functions of extracellular guanosine or eGTP are less studied than adenosine or eATP, their relative concentrations can co-vary, and biological functions of these nucleotides can cross-interact (70,71). GTP is an essential biomolecule that modulates cell signaling via G-proteins and small GTP-binding proteins to facilitate cell proliferation, cell migration, and vesicle trafficking, and, can modulate metabolism and tumor development (72–79). Exocytosis and vesicle secretion can further facilitate release of purinergic nucleotides, inflammatory molecules, enzymes, and ROS into the extracellular milieu, which collectively can alter the TME to become pro-tumorigenic (60,68,80–82) [Figure 5B].

The dose and time-dependence of radiation exposure can significantly alter the impact of RT on tumor microenvironment by affecting tumor or stromal cell behavior, migration, and treatment response (26,28,29,83–90). High-dose irradiation effects include hemorrhage, cognitive decline, neurodegeneration, and premature senescence, which can progress over time (91,92). To identify whether the 20Gy single dose led to aging-like metabolic phenotype or severe neurotoxicity, two aged-mice groups were included (aged (24mo) and aged-obese (24mo)) and their metabolic profiles were compared with 20Gy-IR in C57BL/6 mice cohorts. A hypofractionated treatment cohort 4Gyx10 (total dose=40Gy) was also included to evaluate its correlation with the 20Gy-single dose. Multivariate analysis of ^1^HNMR data revealed a clear distinction between aged-mice groups from 20Gy-single dose [Figure2, Supplementary Figure 2].

The metabolic changes observed in aged and irradiated-mice differed markedly; however, a rise in myo-inositol was observed in both groups. Conversely, age-related markers, scyllo-inositol and sn-glycero-phosphocholine, were only increased in the aged-group, which was accompanied by an overall distinct metabolic signature of these groups in GC-MS [Figure 3, Supplementary Figure 3] (93–95). There were no significant differences in metabolic signatures between 20Gy single dose and 4Gyx10 (total 40Gy) fractionation, indicating the severity of 20Gy single dose-induced damage was nearly equivalent or slightly greater than the more clinically relevant 4Gyx10 fractionated treatment. Increased levels of urea and decreases in NAA and creatine (Cr) or Crn levels are observed in neuropathologies (86,96). We observed a slight increase in urea with radiation in both mouse strains, but NAA and creatinine levels were not consistent and demonstrated a decline only observe in athymic nude mice. These observations indicated a partial neurotoxic state induced by 20Gy-IR; however, it was not as severe as would be expected with higher-dose radiation, and no severe aging-like signatures were observed. This could in part be due to the time-dependence of the experiment, where mice brain samples were harvested for metabolic analysis 24hr after 20Gy-single dose administration, since the tumor injections were performed 24hrs post-IR.

The association between these metabolic effects and time since radiation was investigated by performing a survival study in two mice cohorts, including 1) mice were GBM143-injected 24hrs post-IR to assess immediate or short-term effects (ST-IR) of IR-induced alterations in stroma on tumor development and 2) mice were GBM143-injected 2months post-IR to assess long-term effects (LT-IR) of IR induced stromal alterations on tumor development. Shortest median survival was seen in the LT-IR cohort indicating progressive IR-induced damage in tumor stroma, making it more permissive for tumor growth and recurrence. This corresponds to progressive radiation-induced brain injury and increased susceptibility to progressive neuropathologies in patients treated with RT (97).

## 5. Conclusions

We identified an aggressive tumor behavior and microglial activation following 20Gy single dose brain radiation, which could become more severe with time. Moreover, we found metabolic alterations with a rise in energy carriers (ATP and GTP) and a decline in antioxidants ASC and GSH to associate with the observed tumor phenotype. Since glioblastoma inevitably reoccurs, these observations carry important implications for the impact of the previously radiated microenvironment on the relative aggressiveness of recurrent glioblastoma. Sustained and progressive alterations in the RT-exposed brain microenvironment could worsen GBM outcome.

## 6. Future direction

The role of antioxidants in compromising the therapeutic effect of RT and pro-oxidants in sensitization to RT has long been debated (98–109). Radiation therapy mediates its effects directly or indirectly by production of ROS, thereby causing oxidative damage to macromolecules and induction of apoptosis. Therefore, increased expression of antioxidant peptides in tumors have been thought to reduce the cytotoxic effects of radiotherapy, and GSH inhibition is proposed to have therapeutic advantage in sensitizing cells to RT (59,110). Ascorbate can act as a pro-oxidant in acidic microenvironments, such as tumors (111); thus, it may function as a radio-sensitizer for glioblastoma cells and a radioprotector for normal cells post-RT (112,113). While discrepancies remain regarding ASC’s role as a radio-sensitizer or radio-protector in glioblastoma, its potential as an anticancer agent has been reviewed (114–117).

Our study demonstrates an immediate effect of prior exposure to high-dose irradiation in the non-tumor/untransformed brain cells as decrease in antioxidant levels, including GSH and ASC consistent with their utilization to neutralize RT-induced free radicals. The depletion of these antioxidants can lead to further acute or chronic oxidative stress, altering the brain tumor microenvironment, which may contribute to the enhanced aggressiveness of recurrent tumors. While radiation-induced oxidative stress is necessary for DNA damage in tumor cells, this study raises the question if GSH and ASC administration after completion of radiation could help mitigate the radiation-induced metabolic stress in the microenvironment. If the post-radiation redox state contributes to tumor aggressiveness, there may be an opportunity to attenuate the RT-associated aggressiveness of recurrent GBM, enhancing the long-term safety of brain radiation treatment for glioblastoma.

## Supporting information

Supplementary Excel Sheet 1_NMR

Supplementary Excel Sheet 2_GC-MS

Supplementary Excel Sheet 3_LC-MS

## Acknowledgements

Funding support (TCB) was provided by NIH K12 NRDCP, NINDS NS19770-01, Mayo Clinic Cancer Center, Brains Together for a Cure, the Mayo Clinic Grand Forks Career Development Program and Regenerative Medicine Minnesota. Additionally, this work was supported by the Mayo Clinic Metabolomics Resource Core grant (U24DK100469) and the Mayo Clinic Metabolomics Resource Core NMR developmental funds. The authors acknowledge the editing and research assistance of Superior Medical Experts.

## Author contributions

KG and TB led the project, contributed to experimental design, review, and discussion. TB supervised and supported KG. KG, YX, and BC carried-out mice tumor experiments, KG conducted survival studies. BC supervised KG on irradiator operation. KG performed immunostainings. IO assisted KG. KG, JJ collaborated to analyze images. KG, IV, and SZ performed ^1^H NMR and GC-MS studies, and data analysis. Metabolomics core provided support with LC-MS and data analysis. All authors contributed to experiments and research execution. Figures provided by KG and IV. Illustrations created by KG. All authors contributed to manuscript writing, research, editing and final review.

## Contribution to the field

Radiation therapy (RT) is used as a mainstay treatment modality for cancers, and >50% of all cancer patients receive RT at some stage of their treatment course. RT is the standard of care for glioblastoma has been shown to prolong survival; however, GBM survivors exhibit RT-induced side effects with progressive neuropathological symptoms. Moreover, tumors invariably recur in >90% of all GBM patients. Recent studies have shown damaging side effects caused by RT-exposure of normal tissues surrounding tumors that can cause changes in the brain parenchyma, which can facilitate an aggressive tumor recurrence. Significant advances in radiation therapy have been made to minimize these side-effects, such as image-guided focal radiation treatments or low-fractionated dose administration of total radiation dose (usually of <6Gy/dose fractions) for tumor eradication; however, high-dose radiation treatments are sometimes inevitable during surgical or treatment procedures. Additionally, studies show that late-delayed-effects of low-dose RT treatments can be mimicked by immediate effects of high-dose RT treatments; thus, patients progress to have the same damage as high-dose treatment but in a slow progressive manner over time. Therefore, it is necessary to find alternative post-radiation therapy approaches, which can alleviate tissue intrinsic changes that continue to otherwise progress after RT, leading to late pathological symptoms permitting tumor recurrence. Our work shows that an immediate effect of high-dose IR administration is an increase in energy metabolism in the tissue with decline in levels of tissue antioxidants. Such changes, if left unattended, can fuel the dormant tumor cells that are left after the primary treatment and propagate tumor activation. Thus, administration of antioxidants, such as Glutathione or ascorbate (Vitamin-C) as a post-therapeutic approach can have potential implications in lessening the propensity of tumor recurrence after RT; thereby, also permitting possibilities for use of higher-radiation doses in tumors that are otherwise inaccessible or uncurable by low-fractionated treatment doses.

## Figure Legends

**Supplementary Figure 1: Tumor growth assessed post-moribund for cranially irradiated mice**: A, Scheme for the slicing strategy: Mice brain was sectioned into four equidistant pieces (~1.8mm apart); Slices were made from each in the order of being 5μm thick coronal slices from the cerebral hemisphere only, Rostral to caudal for 22 slides, so as to cover a depth of 120μm from each of the 4 tissue pieces. These slices were arranged onto the glass slides, such that each slice on a slide is obtained from one of the respective 4 brain pieces, sectioned equidistantly. Two slides (1 and 22) were stained with H&E and evaluated for tumor growth. Tumor positive area was detected in slices obtained from two out of 4 sectioned pieces for most of the mice brain samples. Percent positive H and E staining was assessed for each. B, represents the arrangement of the slices on a glass slide, and evaluation of percent positive HE. C, Relative HE staining as observed for slices from 0Gy, 10Gy and 20Gy. Dot-plot for the overall tumor burden estimated in these groups.

**Supplementary Figure 2:** A, (i) Scheme for experiment involving ^1^H-NMR and GC-MS. (ii) Multivariate analysis of ^1^H-NMR data for C57BL/6 mice, having groups as indicated. B, Supervised Orthogonal Projections to Latent Structures Discriminant Analysis, OPLS-DA to show further separation of 0Gy, with irradiated group, IR (20Gy and 4Gyx10); (i) Total VIP (variable importance in the projection) values (ii) Predicted VIP (variable importance in the projection) values. Parameters involved in group separation using multivariate analysis in M1-M7 models are listed in the supplementary Table 1.

**Supplementary Figure 3: Gas chromatography–mass spectrometry (GC-MS)**: The graphs show, relative abundance of metabolites between C57BL/6 mice groups. The significantly altered metabolites are categorized as per their molecular type or biological pathway involvement. Statistical significance is represented as *q ≤ 0.05; **q ≤ 0.01; ***q ≤ 0.001, ****q≤ 0.0001.

**Supplementary Table 1:** Model parameters used for multivariate analysis of ^1^H-NMR data.

