## Supplementary methods for "Radiation Induced Metabolic Alterations Associate with Tumor Aggressiveness and Poor Outcome in Glioblastoma"

Supplementary Material

1. **Materials and Methods**
   1.1 **Liquid chromatography–mass spectrometry (LC-MS)**
       1.1.1 **Microdialysates collected** (300-350 µl) were processed for LC-MS and analyzed on a Quadrupole Time-of-Flight Mass Spectrometer (Agilent Technologies 6550 Q-TOF) coupled with Ultra High-Pressure Liquid Chromatography (1290 Infinity UHPLC Agilent Technologies). Profiling data was acquired under both positive and negative electrospray ionization conditions over a mass range (m/z) of 100–1700 at a resolution of 10,000 (separate runs) in scan mode. Metabolite separation was achieved using two columns of differing  polarity: 1) a hydrophilic interaction column (HILIC, ethylene-bridged hybrid 2.1 x 150 mm, 1.7 mm; Waters) and 2) a reversed-phase C18 column (high-strength silica 2.1 x 150 mm, 1.8 mm; Waters) with gradient described previously (1). A total of four runs per sample were performed to give maximum coverage of metabolites. Samples were injected in duplicate, wherever necessary, and a pooled quality control (QC) sample comprised of all study samples was injected several times during a run. A separate quality control (QC) sample and blank (ringer’s solution) were analyzed with pooled QC to account for analytical and instrumental variability. Dried samples were stored at -20^o^C until analysis. Samples were reconstituted in running buffer and analyzed within 48 hours of reconstitution. Auto-MS/MS data was also acquired with pooled QC sample to aid in unknown compound identification using fragmentation patterns.
       1.1.2 **Data analysis:** Data alignment, filtering, univariate, multivariate statistical and differential analysis was performed using Mass Profiler Professional (Agilent Inc, USA). Metabolites detected in at least ≥80% of one of two groups were selected for differential expression analyses. Metabolite peak intensities and differential regulation of metabolites between groups were determined as described previously (1,2). Each sample was normalized to the internal standard and log2 transformed. Unpaired t-test with multiple testing correction (p<0.05) was used to find the differentially expressed metabolites between two groups.  Default settings were used, except for signal-to-noise ratio threshold (2), mass limit (0.0025 units), and time limit (9s). Putative identification of each metabolite was done based on accurate mass (m/z) against METLIN database (3) using a detection window of ≤7 ppm. The putatively identified metabolites were annotated as Chemical Abstracts Service (CAS), Kyoto Encyclopedia of Genes and Genomes (KEGG), Human Metabolome Project (HMP) database, and LIPID MAPS identifiers. All procedures were performed at the metabolomics core facility, Mayo Clinic, Rochester.
