## Supplementary Figures for "Radiation Induced Metabolic Alterations Associate with Tumor Aggressiveness and Poor Outcome in Glioblastoma"

### Slide 1
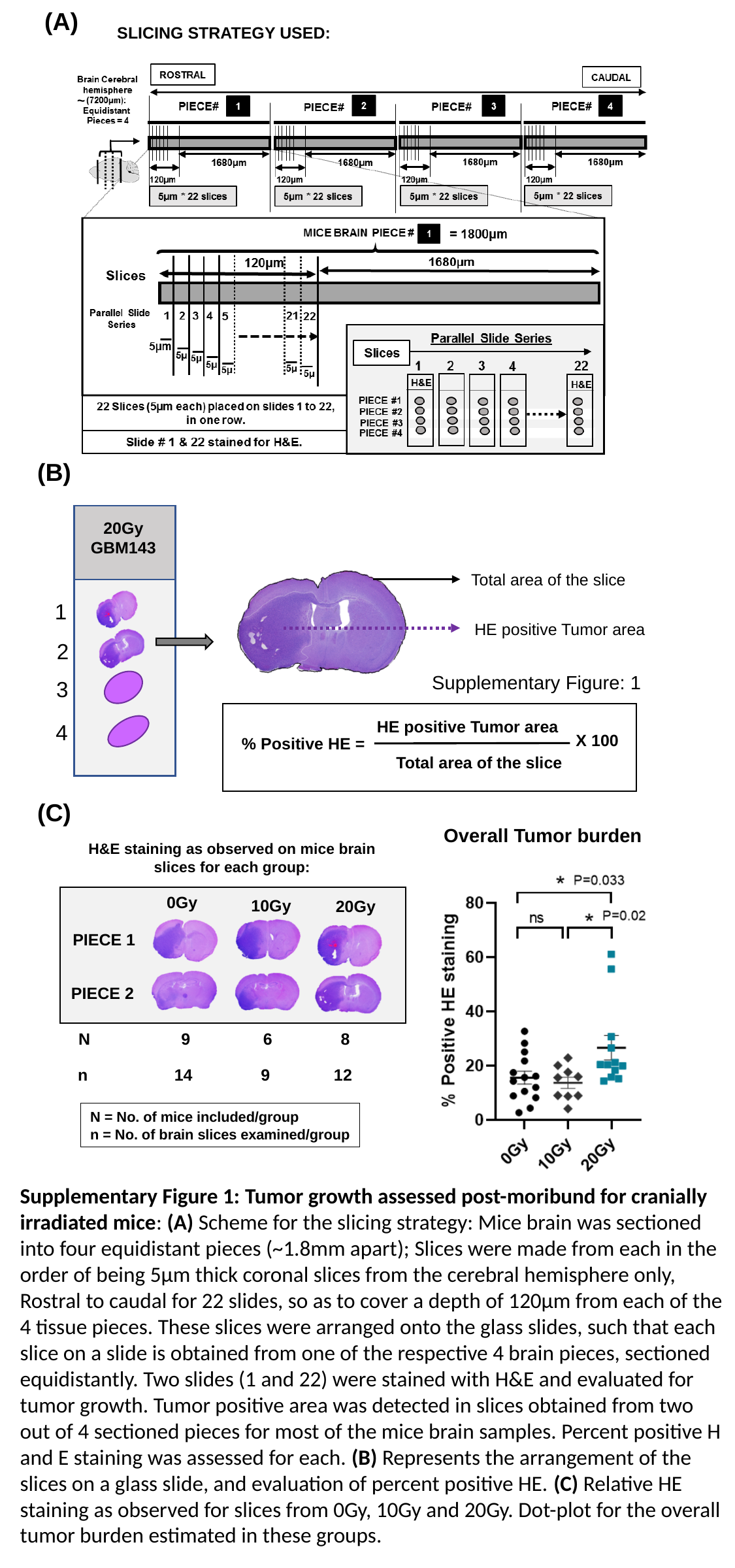

(A)
SLICING STRATEGY USED:
~
(B)
20Gy
GBM143
Total area of the slice
1
HE positive Tumor area
2
Supplementary Figure: 1
3
HE positive Tumor area
X 100
% Positive HE =
Total area of the slice
4
(C)
Overall Tumor burden
H&E staining as observed on mice brain slices for each group:
0Gy
10Gy
20Gy
PIECE 1
PIECE 2
N 9 6 8
n 14 9 12
N = No. of mice included/group
n = No. of brain slices examined/group
Supplementary Figure 1: Tumor growth assessed post-moribund for cranially irradiated mice: (A) Scheme for the slicing strategy: Mice brain was sectioned into four equidistant pieces (~1.8mm apart); Slices were made from each in the order of being 5μm thick coronal slices from the cerebral hemisphere only, Rostral to caudal for 22 slides, so as to cover a depth of 120μm from each of the 4 tissue pieces. These slices were arranged onto the glass slides, such that each slice on a slide is obtained from one of the respective 4 brain pieces, sectioned equidistantly. Two slides (1 and 22) were stained with H&E and evaluated for tumor growth. Tumor positive area was detected in slices obtained from two out of 4 sectioned pieces for most of the mice brain samples. Percent positive H and E staining was assessed for each. (B) Represents the arrangement of the slices on a glass slide, and evaluation of percent positive HE. (C) Relative HE staining as observed for slices from 0Gy, 10Gy and 20Gy. Dot-plot for the overall tumor burden estimated in these groups.

### Slide 2
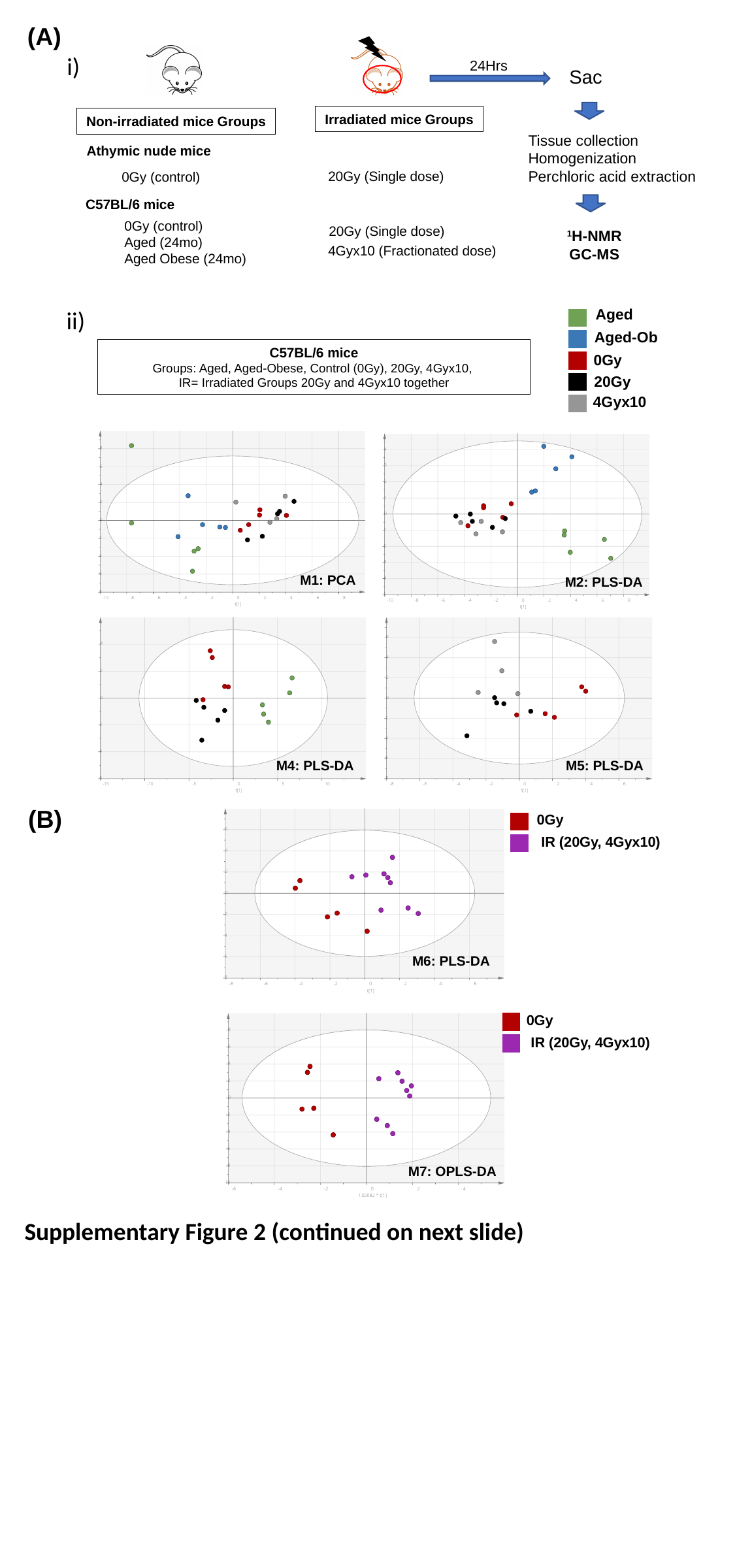

(A)
24Hrs
Sac
Tissue collection
Homogenization
Perchloric acid extraction
Irradiated mice Groups
Non-irradiated mice Groups
Athymic nude mice
20Gy (Single dose)
0Gy (control)
C57BL/6 mice
0Gy (control)
Aged (24mo)
Aged Obese (24mo)
20Gy (Single dose)
4Gyx10 (Fractionated dose)
^1^H-NMR
GC-MS
Aged
Aged-Ob
0Gy
20Gy
4Gyx10
C57BL/6 mice
Groups: Aged, Aged-Obese, Control (0Gy), 20Gy, 4Gyx10,
IR= Irradiated Groups 20Gy and 4Gyx10 together
M2: PLS-DA
M1: PCA
M5: PLS-DA
M4: PLS-DA
M6: PLS-DA
0Gy
IR (20Gy, 4Gyx10)
0Gy
M7: OPLS-DA
IR (20Gy, 4Gyx10)
Supplementary Figure 2 (continued on next slide)
i)
ii)
(B)

### Slide 3
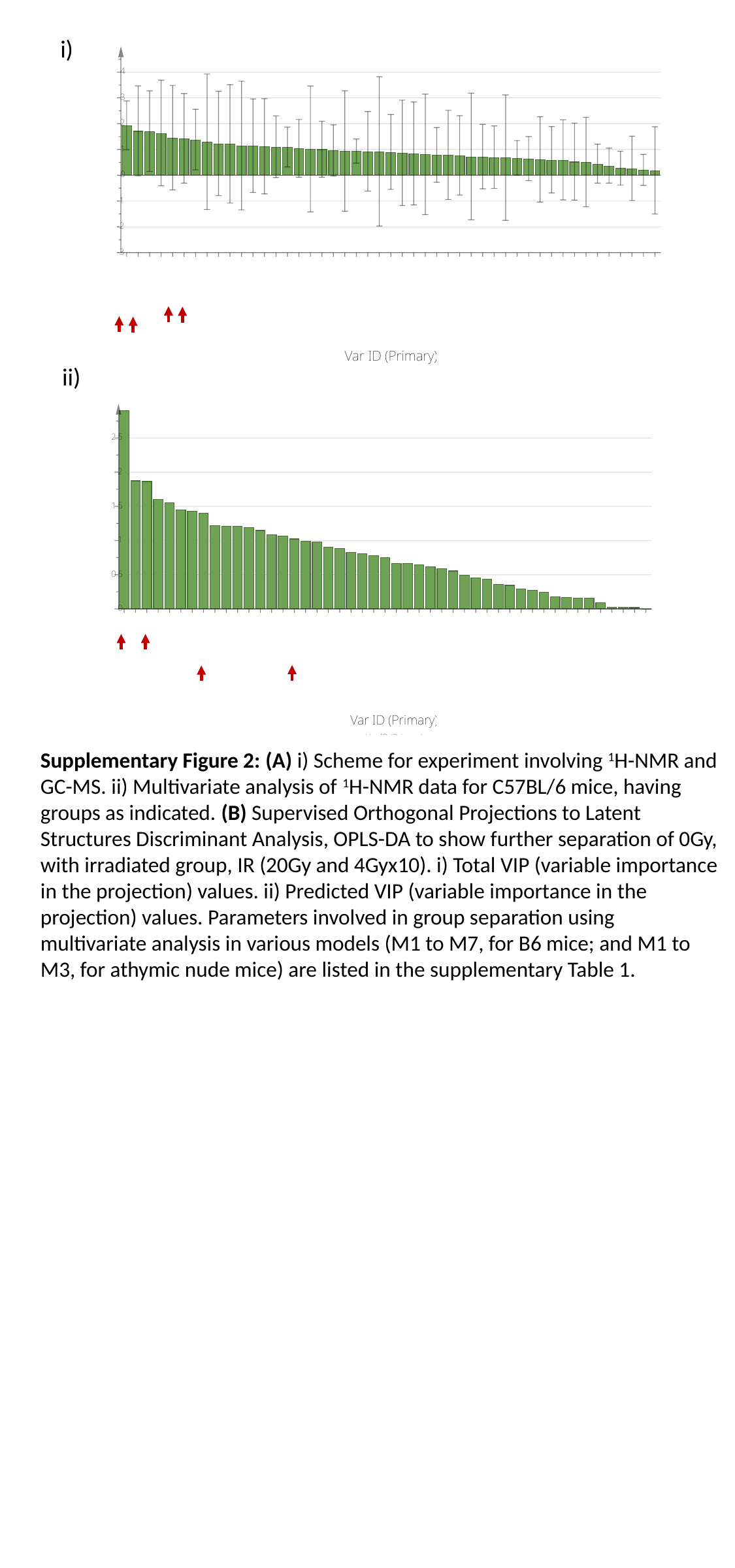

i)
ii)
Supplementary Figure 2: (A) i) Scheme for experiment involving ^1^H-NMR and GC-MS. ii) Multivariate analysis of ^1^H-NMR data for C57BL/6 mice, having groups as indicated. (B) Supervised Orthogonal Projections to Latent Structures Discriminant Analysis, OPLS-DA to show further separation of 0Gy, with irradiated group, IR (20Gy and 4Gyx10). i) Total VIP (variable importance in the projection) values. ii) Predicted VIP (variable importance in the projection) values. Parameters involved in group separation using multivariate analysis in various models (M1 to M7, for B6 mice; and M1 to M3, for athymic nude mice) are listed in the supplementary Table 1.

### Slide 4
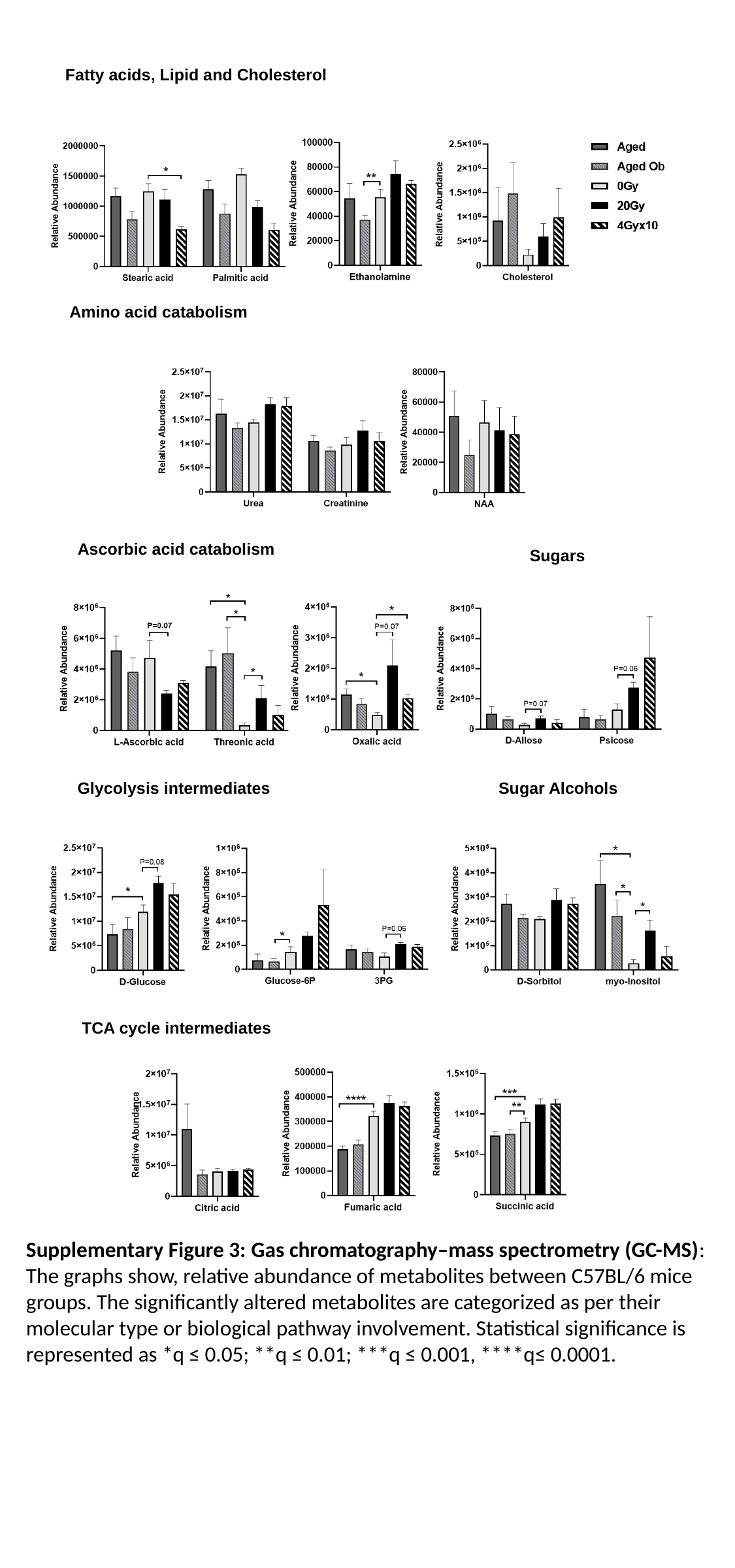

Fatty acids, Lipid and Cholesterol
Amino acid catabolism
Ascorbic acid catabolism
Sugars
Glycolysis intermediates
Sugar Alcohols
TCA cycle intermediates
Supplementary Figure 3: Gas chromatography–mass spectrometry (GC-MS): The graphs show, relative abundance of metabolites between C57BL/6 mice groups. The significantly altered metabolites are categorized as per their molecular type or biological pathway involvement. Statistical significance is represented as *q ≤ 0.05; **q ≤ 0.01; ***q ≤ 0.001, ****q≤ 0.0001.

### Slide 5
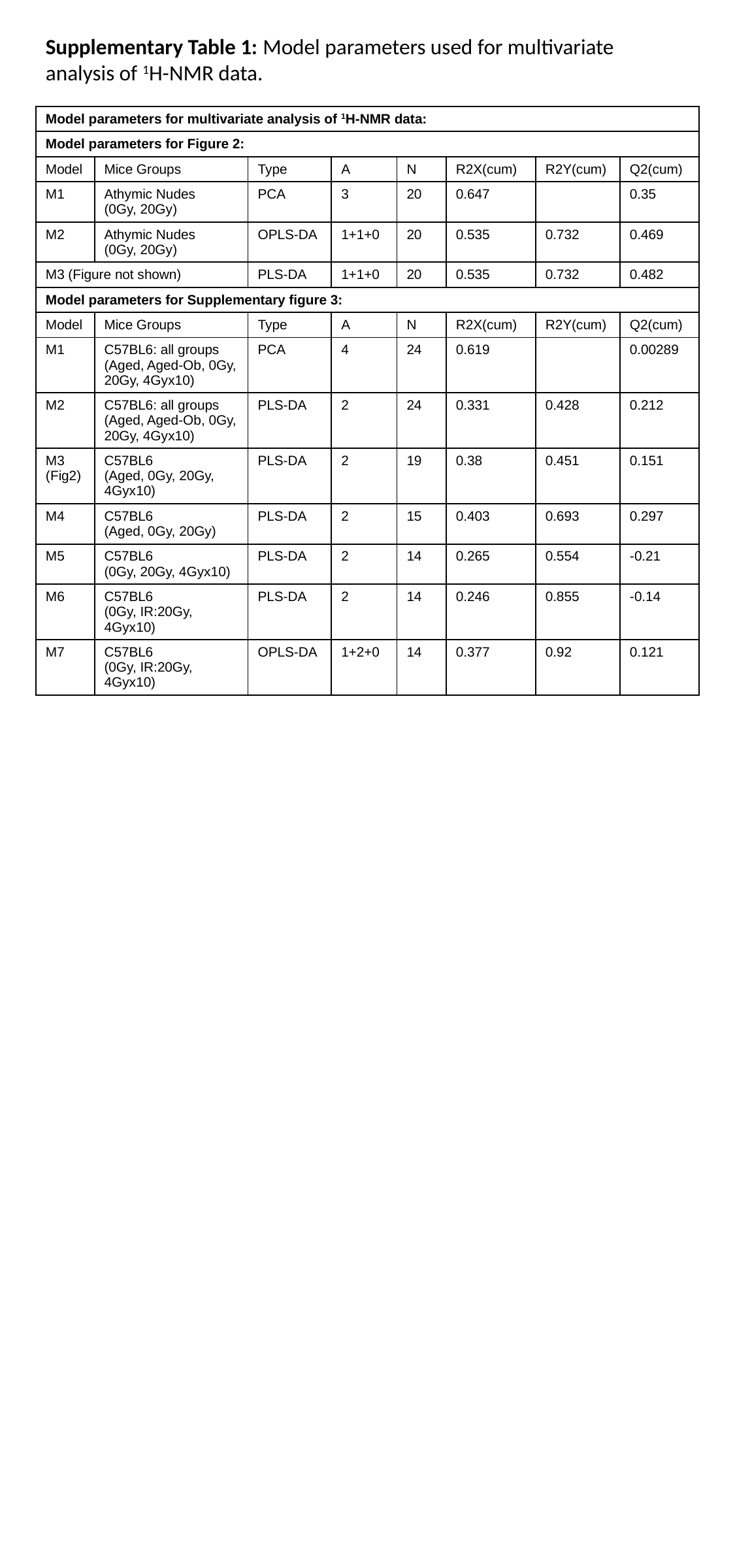

Supplementary Table 1: Model parameters used for multivariate analysis of ^1^H-NMR data.
| Model parameters for multivariate analysis of ^1^H-NMR data: | | | | | | | |
| --- | --- | --- | --- | --- | --- | --- | --- |
| Model parameters for Figure 2: | | | | | | | |
| Model | Mice Groups | Type | A | N | R2X(cum) | R2Y(cum) | Q2(cum) |
| M1 | Athymic Nudes (0Gy, 20Gy) | PCA | 3 | 20 | 0.647 | | 0.35 |
| M2 | Athymic Nudes (0Gy, 20Gy) | OPLS-DA | 1+1+0 | 20 | 0.535 | 0.732 | 0.469 |
| M3 (Figure not shown) | | PLS-DA | 1+1+0 | 20 | 0.535 | 0.732 | 0.482 |
| Model parameters for Supplementary figure 3: | | | | | | | |
| Model | Mice Groups | Type | A | N | R2X(cum) | R2Y(cum) | Q2(cum) |
| M1 | C57BL6: all groups (Aged, Aged-Ob, 0Gy, 20Gy, 4Gyx10) | PCA | 4 | 24 | 0.619 | | 0.00289 |
| M2 | C57BL6: all groups (Aged, Aged-Ob, 0Gy, 20Gy, 4Gyx10) | PLS-DA | 2 | 24 | 0.331 | 0.428 | 0.212 |
| M3 (Fig2) | C57BL6 (Aged, 0Gy, 20Gy, 4Gyx10) | PLS-DA | 2 | 19 | 0.38 | 0.451 | 0.151 |
| M4 | C57BL6 (Aged, 0Gy, 20Gy) | PLS-DA | 2 | 15 | 0.403 | 0.693 | 0.297 |
| M5 | C57BL6 (0Gy, 20Gy, 4Gyx10) | PLS-DA | 2 | 14 | 0.265 | 0.554 | -0.21 |
| M6 | C57BL6 (0Gy, IR:20Gy, 4Gyx10) | PLS-DA | 2 | 14 | 0.246 | 0.855 | -0.14 |
| M7 | C57BL6 (0Gy, IR:20Gy, 4Gyx10) | OPLS-DA | 1+2+0 | 14 | 0.377 | 0.92 | 0.121 |
